# A practical guide for improving transparency and reproducibility in neuroimaging research

**DOI:** 10.1101/039354

**Authors:** Krzysztof J. Gorgolewski, Russell A. Poldrack

## Abstract

Recent years have seen an increase in alarming signals regarding the lack of replicability in neuroscience, psychology, and other related fields. To avoid a widespread crisis in neuroimaging research and consequent loss of credibility in the public eye, we need to improve how we do science. This article aims to be a practical guide for researchers at any stage of their careers that will help them make their research more reproducible and transparent while minimizing the additional effort that this might require. The guide covers three major topics in open science (data, code, and publications) and offers practical advice as well as highlighting advantages of adopting more open research practices that go beyond improved transparency and reproducibility.

## Introduction

The question of how the brain creates the mind has captivated humankind for thousands of years. With recent advances in human *in vivo* brain imaging, we how have effective tools to peek into biological underpinnings of mind and behavior. Even though we are no longer constrained just to philosophical thought experiments and behavioral observations (which undoubtedly are extremely useful), the question at hand has not gotten any easier. These powerful new tools have largely demonstrated just how complex the biological bases of behavior actually are. Neuroimaging allows us to give more biologically grounded answers to burning questions about everyday human behavior (“why do we crave things?”, “how do we control learned responses?”, “how do we regulate emotions?” etc.), as well as influencing how we think about mental illnesses.

In addition to fantastic advances in terms of hardware we can use to study the human brain (function Magnetic Resonance Imaging, Magnetoencephalography, Electroencephalography etc.) we have also witnessed many new developments in terms of data processing and modelling. Many bright minds have contributed to a growing library of methods that derive different features from brain signals. Those methods have widened our perspective on brain processes, but also resulted in methodological plurality [1]. Saying that there is no single best way to analyze a neuroimaging dataset is an understatement; we can confidently say that there many thousands of ways to do that.

Having access to a plethora of denoising and modelling algorithms can be both good and bad. On one side there are many aspects of brain anatomy and function that we can extract and use as dependent variables, which maximizes the chances of finding the most appropriate and powerful measure to ask a particular question. On the other side, the incentive structure of the current scientific enterprise combined with methodological plurality can be a dangerous mix.

Scientists rarely approach a problem without a theory, hypothesis, or a set of assumptions, and the high number of “researcher degrees of freedom” [2] can implicitly drive researchers to choose analysis workflows that provide results that are most consistent with their hypotheses.

As Richard Feynman said “The first principle is that you must not fool yourself — and you are the easiest person to fool.”. Additionally, neuroimaging (like almost every other scientific field) suffers from publication bias, in which “null” results are rarely published, leading to overestimated effect sizes (for review of this and other biases see [3].

Recent years have seen an increase in alarming signals about the lack of replicability in neuroscience, psychology, and other related fields [4]. Neuroimaging studies generally have low statistical power (estimated at 8%) due to the high cost of data collection which results in an inflation of the number of positive results that are false [5],. To avoid a widespread crisis in our field and consequently losing credibility in the public eye, we need to improve how we do science. This article aims to complement existing literature on the topic [6–8] by compiling a practical guide for researchers at any stage of their careers that will help them make their research more reproducible and transparent while minimizing the additional effort that this might require.

**Figure 1.**
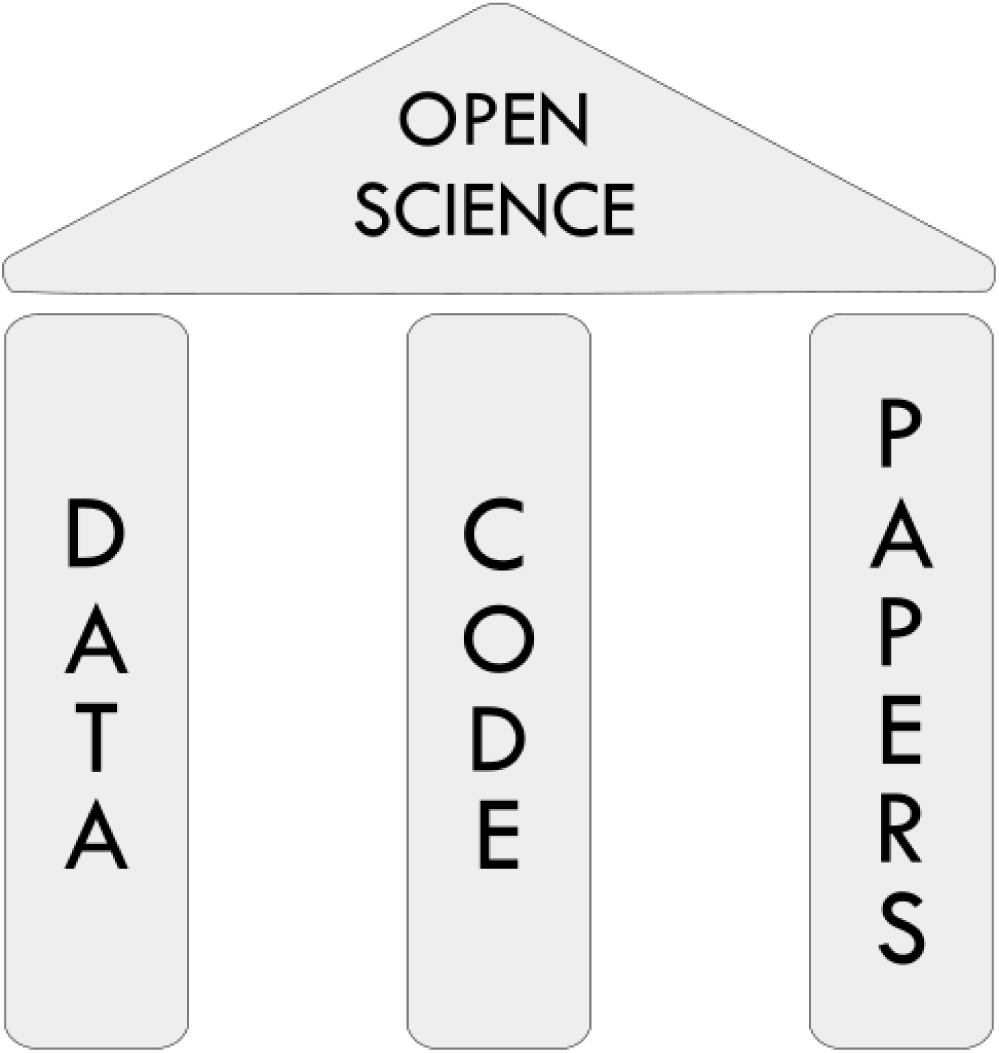
Three pillars of Open Science: data, code, and papers.

## How to deal with data

Data are a central component of the scientific process. When data are made open accessible, they not only allow the scientific community to validate the accuracy of published findings, but also empower researchers to perform novel analyses or combine data from multiple sources. Papers accompanied by publicly available data are on average cited more often [9,10], while at the same time exposing fewer statistical errors [11]. Data sharing has been mandated by some grant funding agencies, as well as journals. Some also argue that sharing data is an ethical obligation toward study participants, in order to maximize the benefits of their participation [12]. Neuroimaging has a substantial advantage in terms of ease of data capture since the data generation process is completely digital. In principle one could provide a digital record of the entire research process for the purpose of reproducibility. However, even though data sharing in neuroimaging has been extensively reviewed in [13] and [14] there is little practical advice on the topic.

### Consent forms

Planning for data sharing should start at the ethical approval stage. Even though in the United States de-identified data can be freely shared without specific participant consent, the rules differ in other countries (and they may change in the upcoming revisions to the Common Rule, which governs research in the US). In addition it is only fair to inform your participants about your intention to maximize their generous gift by sharing their data, and to allow them to withdraw from research if they don’t wish to have their data shared. However, consent form language needs to be carefully crafted. To streamline the creation of consent forms with data sharing clauses, we have prepared a set of templates that can be easily inserted into existing consent forms after minor adjustments^1^. Those templates have been derived from existing consent forms of leading data sharing projects (Nathan Kline Institute Enhanced sample [15] and Human Connectome Project [16]) followed by consultations with bioethics experts. The templates come in two flavors: one for normal populations and generic data and one for sensitive populations and/or data. The latter splits the data into two sets: a publicly available portion and a portion that requires approval of a data sharing committee (that would assess the ability of the applicant to protect sensitive data) in order to gain access to. We recommend using the restricted access version only for data and populations for which a) potential data re-identification is easy due to small sample and/or the level of detail of included variables (for example exact time and location of scanning) or b) re-identification would lead to negative consequences for the participants (for example in a study of HIV-positive subjects).

### Data organization

To successfully share data one has to properly describe it and organize it. Even though some experimental details such as the MRI phase encoding direction may seem obvious for the researcher who obtained the data, they need to be clearly explained for external researchers. In addition, good data organization and description can reduce mistakes in analysis. While each experiment is different and may include unique measurements or procedures, most MRI datasets can be accurately described using one fairly simple scheme. Recently we have proposed such scheme - the Brain Imaging Data Structure (BIDS) [17]. It was inspired by the data organization used by OpenfMRI database, but has evolved through extensive consultations with the neuroimaging community. BIDS aims at being simple to adopt,and roughly follows existing practices common in the neuroimaging community. It is heavily based on a specific organization of files and folders and uses simple file formats such as NifTI, tab-separated text and JSON. It does not require a database or any external piece of software for processing. A browser-based validator has been developed that allows one to easily check whether a dataset accurately follows the BIDS standard^2^.

An additional benefit of using a standardized data organization scheme is that it greatly streamlines the data curation that is necessary when submitting data to a data sharing repository. For example datasets formatted according to BIDS undergo a faster and more streamlined curation process when submitted to OpenfMRI database [18].

### Publishing data

Data should be submitted to a repository before submitting the relevant paper. This allows the author to point the readers and reviewers to the location of the data in the manuscript. The manuscript can benefit from increased transparency due to shared data and the data itself can become a resource enabling additional future research.

The most appropriate places for depositing data are field-specific repositories. Currently in human neuroimaging there are two well recognized repositories accepting data from everyone: FCP/INDI [19] (for any datasets that include resting state fMRI and T1 weighted scans) and OpenfMRI [18] (for any datasets that include any MRI data). Field specific repositories have the advantage of more focused curation process that can greatly improve the value of your data. They also increase data discoverability since researchers search through them first when looking for datasets, and some (like OpenfMRI) are indexed by PubMed which allows the dataset to be directly linked to the paper via the LinkOut mechanism.

If for some reason field specific repositories are not an option we recommend using field agnostic repositories such as FigShare, Dryad, or DataVerse. When picking a repository one should think of long term data retention. No one can guarantee existence of a repository in the far future, but historical track record and the support of well established institutions can increase the chances that the data will be available in the decades to come. In addition a platform such as Open Science Framework (www.osf.io) can be used to link together datasets deposited in field agnostic repositories with code and preprints (see below). If one is concerned about losing competitive advantage by sharing data before the relevant manuscript will be accepted and published (so called “scooping”) one can consider setting an embargo period on the submitted dataset. OSF^3^, figshare^4^, and Dryad^5^ support this functionality.

Since data-agnostic repositories do not impose any restriction on the form in which you deposit your data nor do they check completeness, it is essential to ensure that all of the necessary data and metadata are present. Using a data organization scheme designed for neuroimaging needs such as BIDS or XCEDE [20] can help ensure that data are represented accurately. In addition, it is a good idea to ask a colleague who is unfamiliar with the data to evaluate the quality and completeness of the description.

If the data accompanying the paper is very large or particularly complex you should consider writing a separate data paper to describe the dataset [21]. A data paper is new type of publication dedicated purely to description of the data rather than its analysis. It can provide more space to describe the experimental procedures and data organization details, and also provides a mechanism for credit when the data are reused in the future. In addition, one often receives useful feedback about the dataset description through the peer review process. The list of journals that currently accept neuroimaging data papers includes but is not limited to: Scientific Data, Gigascience, Data in Brief, F1000Research, Neuroinformatics, and Frontiers in Neuroscience.

In addition to raw data we also encourage authors to share derivatives such as preprocessed volumes, statistical maps or tables of summary measures. Because other researchers are often interested in reusing the results rather than the raw data, this can further increase the impact of the data. For example, statistical maps can be used to perform image-based meta analysis or derive regions of interest for new studies. For sharing statistical maps we encourage authors to use the NeuroVault.org platform [22]. The UCLA Multimodal Connectivity Database [23] provides similar service but for connectivity matrices (derived from fMRI or DWI data).

Finally published data should be accompanied by an appropriate license. Data are treated differently by the legal system than creative works (i.e. papers, figures) and software and thus require special licenses. Following the lead of major scientific institutions such as BioMed Central, CERN, or The British Library we recommend using an unrestricted Public Domain license (such as CC0 or PDDL) for data^6^. Using such license would maximize the impact of the shared data, by not imposing any restriction on how it can be used and combined with other data. The appropriate legal language that needs to accompany your data can be obtained from https://creativecommons.org/publicdomain/zero/1.0 or http://opendatacommons.org/licenses/pddl//. There are also other more restrictive license options (see http://www.dcc.ac.uk/resources/how-guides/license-research-data). However, additional restrictions can have unintended consequences. For example, including a Non-Commercial clause, while seemingly innocuous, could in its broadest interpretation prevent your data from being used for teaching or research at a private university. Similarly, a No-Derivatives clause can prevent your data from being combined in any form with other data (for example a brain template released under No-Derivatives license cannot be used as a coregistration target).

## How to deal with code

Neuroimaging data analysis has required computers since its inception. A combination of compiled or script code is involved in every PET, MRI, or EEG study, as in most other fields of science. The code we write to analyze data is a vital part of the scientific process, and similar to data, is not only necessary to interpret and validate results, but can be also used to address new research questions. Therefore the sharing of code is as important as the sharing of data for scientific transparency and reproducibility.

Because most researchers are not trained in software engineering, the code that is written to analyze neuroimaging data (as in other areas of science) is often undocumented and lacks the formal tests that professional programmers use to ensure accuracy. In addition to the lack of training, there are few incentives to spend the time necessary to generate high-quality and well-documented code. Changes in the incentive structure of science will take years, but in the meantime, perceived poor quality of code and lack of thorough documentation should not prevent scientists from publishing it [24]. Sharing undocumented code is a much better than not sharing code at all and can still provide benefits to the author. Perhaps the most compelling motivation for sharing code comes from citation rates. Papers accompanied by usable code are on average cited more often than their counterparts without the code [25].

An additional concern that stops researchers from sharing code is fear that they will have to provide user support and answer a flood of emails from other researchers who may have problems understanding the codebase. However, sharing code does not oblige a researcher to provide user support. One useful solution to this problem is to set up a mailing list (for example with Google) and point all users to ask questions through it; in this way, answers are searchable, so that future users with the same questions can find them via a web search. Alternatively one can point user to a community driven user support forum for neuroinformatics (such as NeuroStars.org) and ask them to tag their questions with a label uniquely identifying the software or script in question; we have found this to be a useful support solution for the OpenfMRI project. Both solutions foster a community that can lead to users helping each other with problems, thus relieving some of the burden from the author of the software. In addition, since the user support happens through a dedicated platform there is less pressure on the author to immediately address issues than there would be with user requests send directly by email.

Many of the issues with code quality and ease of sharing can be addressed by careful planning. One tool that all research programmers should incorporate into their toolbox is the use of a Version Control System (VCS) such as git. VCS provides a mechanism for taking snapshots of evolving codebase that allow tracking of changes and reverting them if there is a need (e.g., after making a change that ends up breaking things). Adopting a VCS leads a to cleaner code base that is not cluttered by manual copies of different versions of a particular script (e.g, “script_version3_good_Jan31_try3.py”). VCS also allows one to quickly switch between branches - alternative and parallel versions of the codebase - to test a new approach or method without having to alter a tried and tested codebase. For a great introduction to git we refer the reader to [26]. We encourage scientists to use git rather than other VCS due to a passionate and rapidly growing community of scientists who use the GitHub.com platform, which is a freely available implementation of the git VCS system. In the simplest use case GitHub is a platform for sharing code (which is extremely simple for those who already use git as their VCS), but it also includes other features which make contributing to collaborative projects, reviewing, and testing code simple and efficient. The Open Science Framework mentioned above can be used to link together data and code related to a single project. It can also be used to set embargo period on the code so it could be submitted with the paper while minimising the risk of “scooping”.

Striving for automation whenever possible is another strategy that will not only result in more reproducible research, but can also save a lot of time. Some analysis steps seem to be easy to perform manually, but that remains true only when they need to be performed just once. Quite often in the course of a project parameters are modified, list of subjects are changed, and processing steps need to be rerun. This is a situation in which having a set of scripts that can perform all of the processing steps automatically instead of relying on manual interventions can really pay off. There are many frameworks that help design and efficiently run neuroimaging analyses in automated fashion. Those include, but are not limited to: Nipype [27], PSOM [28], aa [29], and make [30]. As an example, for our recent work on the MyConnectome project[31] we created a fully automated analysis pipeline, which we implemented using a virtual machine^7^.

While automation can be very useful for reproducibility, the scientific process often involves interactive interrogation of data interleaved with notes and plots. Fortunately there is a growing set of tools that facilitate this interactive style of work while preserving a trace of all the computational steps, which increases reproducibility. This philosophy is also known as “literate programming” [32] and combines analysis code, plots, and text narrative. The list of tools supporting this style of work includes, but is not limited to: Jupyter (for R, Python and Julia)^8^, R Markdown (for R)^9^ and matlabweb (for MATLAB)^10^. Using one of those tools not only provides the ability to revisit an interactive analysis performed in the past, but also to share an analysis accompanied by plots and narrative text with collaborators. Files created by one of such systems (in case of Jupyter they are called Notebooks) can be shared together with the rest of the code on GitHub, which will automatically render included plots so they can be viewed directly from the browser without requiring installation of any additional software.

As with data, it is important to accompany shared code with an appropriate license. Following [6] we recommend choosing a license that is compatible with the open source definition such as Apache 2.0, MIT, or GNU General Public License (GPL)^11^. The most important concept to understand when choosing a license is “copyleft”. A license with a “copyleft” property (such as GPL) allows derivatives of your software to be published, but only if done under the same license. This property limits the range of code your software can be combined with (due to license incompatibility) and thus can restrict the reusability of your code; for this reason, we generally employ minimally restrictive licenses such as the MIT license. Choosing an open source license and applying it to your code can be greatly simplified by using a service such as choosealicense.com.

## How to deal with publications

Finally, the most important step in dissemination of results is publishing a paper. An essential key to increasing transparency and reproducibility of scientific outputs is accurate description of methods and data. This not only means that the manuscript should include links to data and code mentioned before (which entails that both data and code should be deposited before submitting the manuscript), but also thorough and detailed description of methods used to come to a given conclusion. As an author one often struggles with a fine balance between detailed description of different analyses performed during the project and and the need to explain the scientific finding in the most clear way. It is not unheard of that for the sake of a better narrative some results are omitted^12^. At the same time there is a clear need to present results in a coherent narrative with a clear interpretation that binds the new results with an existing pool of knowledge^13^. We submit that one does not have exclude the other. A clear narrative can be provided in the main body of the manuscript and the details of methods used together with null results and other analyses performed on the dataset can be included in the supplementary materials, as well as in the documentation of the shared code. In this way, the main narrative of the paper is not obfuscated too many details and auxiliary analyses, but all of the results (even null ones) are available for the interested parties. Such results from extra analyses could include for example all of the additional contrasts that were not significant and thus not reported in the main body of the manuscript (of which unthresholded statistical maps should be shared for example using a platform such as NeuroVault). Often these extra analyses and null results may seem uninteresting from the author's point of view, but one cannot truly predict what other scientists can be interested in. In particular, the null results (which are difficult to publish independently) can contribute to growing body of evidence that can be used in the future to perform meta analyses.For more extensive set of recommendation for reporting neuroimaging studies, see the recent report from the Organization for Human Brain Mapping’s Committee on Best Practices in Data Analysis and Sharing (COBIDAS) report^14^.

The last important topic to cover is accessibility of the manuscript. To maximize the impact of published research one should consider making the manuscript publicly available. In fact many funding bodies (NIH, Wellcome Trust) require this for all manuscripts describing research that they have funded. Many journals provide an option to make papers open access, albeit sometimes at prohibitively high price (for example the leading specialist neuroimaging journal - NeuroImage - requires a fee of $3000). Unfortunately the most prestigious journals (Nature and Science) do not provide such option despite many requests from the scientific community. Papers published in those journals remain “paywalled” - available only through institutions which pay subscription fees, or through public repositories (such as PubMed Central) after a sometimes lengthy embargo period. The scientific publishing landscape is changing [33,34], and we hope it will evolve in a way that will give everyone access to published work as well as to the means of publication. In the meantime we recommend ensuring open access by publishing preprints at Bioxiv or arXiv before submitting the paper to a designated journal. In addition to making the manuscript publicly available without any cost, this solution has other advantages. Firstly it allows the wider community to give feedback to the authors about the manuscript and potentially improve it which is beneficial for both the authors as well as the journal the paper will be submitted to; for example, the present paper received useful comments from three individuals in addition to the appointed peer reviewers. Secondly, in case of hot topics publishing a preprint establishes precedence on being the first one to describe a particular finding. Finally since preprints have assigned DOIs other researchers can reference them even before they will be published in a journal. Preprints are increasingly popular and vast majority of journals accept manuscripts that have been previously published as preprints. We are not aware of any neuroscience journals that do not allow authors to deposit preprints before submission, although some journals such as *Neuron* and *Current Biology* consider each submission independently and thus one should contact the editor prior to submission.

To further improve accessibility and impact of research outputs one can also consider sharing papers that have already been published in subscription based journals. Unfortunately this can be difficult due to copyright transfer agreements many journals require from authors. Such agreement give the journal exclusive right to the content of the paper. However, each publisher uses a different set of rules and some of them allow limited sharing of your work you have surrender your rights to. For example Elsevier (publisher of Neuroimage) allows authors to publish their accepted manuscripts (without the journal formatting) on a non-commercial website, a blog or a preprint repository^15^. Wiley (publisher of Human Brain Mapping) has a similar policy for submitted manuscripts (before the paper gets accepted), but requires an embargo of 12 months before authors can share the accepted manuscript^16^. Policies for other journals might vary. SherPa/ROMEO (http://www.sherpa.ac.uk/romeo) is a databaset that allows authors to quickly check what the journal they published with allows to share and when.

There are multiple options when it comes to choosing a repository to share manuscripts published in subscription-based journals. Private websites, institutional repositories, and preprint servers seems to be well within the legal restrictions of most journals. Commercial websites such as researchgate.com and academia.edu remain a legal grey zone (with some reports of Elsevier taking legal actions to remove papers from one of them^17^). If the research has been at least partially funded by NIH one can deposit the manuscript in PubMed Central (respecting appropriate embargos)^18^.

## Discussion

In this guide we have carefully selected a list of enhancements that every neuroimaging researcher can make to their scientific workflow that will improve the impact of their research, benefiting not only them individually the community as a whole. We have limited the list to mechanisms that have been tested and discussed in the community for number of years and which have clear benefits to the individual researcher. However, the way science is conducted is evolving constantly and there are many more visions that could be implemented. In the following section we discuss some of the emerging trends that may become commonplace in the future.

### Pre-registration

We have mentioned in the introduction that the field of neuroimaging is both blessed and cursed with plurality of analysis choices which can lead to biases in published results (since many decisions about statistical treatment of data are made after seeing the data). We recommended taking advantage of supplementary materials to elaborate on all performed analyses and sharing statistical maps of null effect contrasts as a partial remedy of this problem. However, further reduction of publication bias can be achieved even more effectively by adopting the pre-registration mechanism [35]. This way of doing research, originally adopted from clinical trials, involves writing and registering (in a third party repository) a study plan outlining details of data acquisition, subject exclusion criteria, and planned analyses even before that data have been acquired. This not only motivates researchers to formulate hypotheses before seeing data, but also allows for a clear distinction between results of hypothesis driven confirmatory analyses (included in the pre-registration) and exploratory analyses (added after seeing the data). It is worth mentioning that exploratory analyses are by no means inferior to confirmatory analyses; they are an important part of science, generating new hypotheses that can be tested by future studies. However exploratory analyses can suffer from bias (since their inception was influenced by the data itself) and thus require additional evidence. Unfortunately, confirmatory and exploratory analyses are often not properly distinguished in publications, a problem that could be remedied by preregistration. Preregistration also plays a vital role in highlighting hypotheses that turned out not to be confirmed by the data (“null effects”).

It is clear that preregistration can help in research transparency and reproducibility by reducing biases. It is also important to acknowledge that putting together and registering a binding research plan requires a significant time investment from the researcher and thus is not common a common practice (with exception to replication studies [4,36]). There are, however, additional incentives for individual researchers to preregister their studies. For example, the Center for Open Science spearheaded the Registered Reports^19^ initiative in 2012. According to this mechanism, authors send their preregistration reports (Introduction, Methods parts of a future paper and optionally analysis of pilot data) for peer review to a journal for peer review. Validity of the experimental plan is assessed and if deemed sufficient receives “In-principle acceptance” (IPA), in which case the journal guarantees to publish the final version of the paper (after data collection and analysis) independently of the results (i.e. even if the hypothesized effect was not found). Currently journals accepting in neuroimaging papers participating in the Registered Reports program include: AIMS Neuroscience [37], Attention, Perception, and Psychophysics, Cognition and Emotion [38], Cortex [39] and European Journal of Neuroscience. Additionally, The Center for Open Science started a Preregistration Challenge^20^ providing $1000 reward for the first 1000 preregistered eligible studies. This initiative is independent of the Registered Reports and does not guarantee publication, but the list of eligible journals is much longer (includes such journals as PloS Biology, Hippocampus, or Stroke).

### Peer review and giving feedback

An important part of the scientific method is peer review but with a few notable exceptions (eLife, GigaScience, ScienceOpen, and F1000Research), the review procedure happens behind closed doors and thus leaves the reader without any information on how a published paper was evaluated (other than the fact that it was accepted). In addition, at most journals reviewers do not get credit for their hard work, though some (such as the Frontiers journals) list the reviewers on each published paper. This situation can be remedied by publishing reviews performed for journals after the paper has been published. Several outlets exists that allow that. PubMed Commons allow registered and verified users of PubMed to provide comments under every paper indexed by PubMed. Those comments have to be signed so there is no option to remain anonymous (which is important for junior researchers afraid of a blowback after criticizing work from an established lab). Another option is PubPeer - a website that allow anyone to comment on any published paper or preprint. It supports both anonymous and signed comments so it’s up to the reviewer to decide what is better for them. Finally there is Publons.com - a platform for tracking reviewers profiles and publishing reviews. Thanks to collaborations with many journals it is very easy to use and even allows you to get credit for publishing your reviews anonymously.

All of those platforms can be used not only to share reviews solicited from reviewers by journals, but also to share comments and give feedback about already published work or preprints shared by other researchers. Peer review expanded to the whole community can improve the quality of research, catch mistakes, or help with the clarity of both preprints and already published work. Giving feedback on preprints can be especially useful when it comes to highlighting already published work that authors might have missed (which considering the number of papers published every year is not unlikely).

Signing openly shared reviews can have some benefits when it comes to establishing one's reputation as an expert in the field. Well thought through and carefully worded reviews consisting of constructive criticism are hard to come by and extremely valuable. By sharing and signing reviews researchers can not only help their peers, but also boost their reputation which can potentially seen favourably by hiring committees and grant review boards. However, we feel that the option of anonymous reviews remains very important since on many occasions it will be the only way for researchers to express concerns about validity of some work.

##### Box 1. Simple steps towards open science

Data:

- Include a section about data sharing to your consent forms.
- Share your raw data upon paper submission using a repository dedicated for neuroimaging.
- Consider writing a separate data paper for more complex and interesting datasets.
- Remember that sharing your data improves the impact and citation rates of your research!
Code:

- Use version control system for all your projects.
- Share your code on GitHub.com even if it’s not well documented.
- Set up a mailing list for user related questions.
- People reusing the code you shared will cite the relevant papers.
Papers:

- Include all extra analyses and null results in the supplementary materials without sacrificing the clarity of the message in the main body of the manuscript.
- Submit preprints to claim precedence, solicit feedback and give access to your research.

## Summary

The scientific method is evolving towards a more transparent and collaborative endeavour. The age of digital communication allows us to go beyond printed summaries and dive deeper into underlying data and code. In this guide we hope to have shown that there are many improvements in scientific practice everyone can implement with relatively little added effort that will improve transparency, replicability and impact of their research. Even though the added transparency might in rare cases expose errors those are a natural part of the scientific process, as a community researchers should acknowledge their existence and try to learn from them instead of hiding them and antagonizing those who make them.

## Acknowledgements

This work was supported by Laura and John Arnold Foundation, National Science Foundation, and National Institutes of Health. We would also like to thank our reviewers: Nikolas Kriegeskorte, Brian Nosek, Cyril Pernet, Yaroslav Halchenko, and Cameron Craddock for helpful comments and corrections as well as Michael Milham for stimulating discussion.

## Footnotes

1 https://openbrainconsent.readthedocs.org/en/latest/ultimate.html

2 http://incf.github.io/bidsvalidator

3 https://osf.io/faq/

4 https://figshare.com/blog/The_future_of_figshare/166

5 http://datadryad.org/pages/faq

6 https://wiki.creativecommons.org/wiki/CC0_use_for_data

7 https://github.com/poldrack/myconnectomevm

8 http://jupyter.org

9 http://rmarkdown.rstudio.com

10 https://www.ctan.org/pkg/matlabweb

11 https://opensource.org/licenses

12 http://sometimesimwrong.typepad.com/wrong/2015/11/guestpostataleoftwopapers.html

13 http://www.russpoldrack.org/2015/11/aregoodscienceandgreatstorytelling.html

14 www.humanbrainmapping.org/cobidas/

15 https://www.elsevier.com/about/companyinformation/policies/sharing

16 http://olabout.wiley.com/WileyCDA/Section/id826716.html

17 http://svpow.com/2013/12/06/elsevieristakingdownpapersfromacademiaedu/

18 https://nihms.nih.gov

19 https://osf.io/8mpji

20 https://cos.io/prereg/

